# Temporal transcriptomic dynamics in developing macaque neocortex

**DOI:** 10.1101/2023.07.13.548828

**Authors:** Longjiang Xu, Zan Yuan, Jiafeng Zhou, Yuan Zhao, Wei Liu, Shuaiyao Lu, Zhanlong He, Boqin Qiang, Pengcheng Shu, Yang Chen, Xiaozhong Peng

## Abstract

Despite intense research on mice, the transcriptional regulation of neocortical neurogenesis remains limited in humans and non-human primates. Cortical development in rhesus macaque is known to recapitulate multiple facets of cortical development in humans, including the complex composition of neural stem cells and the thicker supragranular layer. To characterize temporal shifts in transcriptomic programming responsible for differentiation from stem cells to neurons, we sampled parietal lobes of rhesus macaque at E40, E50, E70, E80, and E90, spanning the full period of prenatal neurogenesis. Single-cell RNA sequencing produced a transcriptomic atlas of the developing rhesus macaque neocortex parietal lobe. Identification of distinct cell types and neural stem cells emerging in different developmental stages revealed a terminally bifurcating trajectory from stem cells to neurons. Notably, deep-layer neurons appear in the early stages of neurogenesis, while upper-layer neurons appear later. While these different lineages show overlap in their differentiation program, cell fates are determined post-mitotically. Pseudotime trajectories from vRGs to oRGs revealed differences in dynamic gene expression profiles and identified divergence in their activation of BMP, FGF, and WNT signaling pathways. These results provide a comprehensive picture of the temporal patterns of gene expression leading to different fates of radial glial progenitors during neocortex layer formation.

## Introduction

The neocortex is the center for higher brain functions, such as perception and decision-making. Therefore, the dissection of its developmental processes can be informative of the mechanisms responsible for these functions. Several studies have advanced our understanding of the neocortical development principles in different species, especially in mice. Generally, the dorsal neocortex can be anatomically divided into six layers of cells occupied by distinct neuronal cell types. The deep-layer neurons project to the thalamus (layer VI neurons) and subcortical areas (layer V neurons), while neurons occupying more superficial layers (upper-layer neurons) preferentially form intracortical projections^1^. The generation of distinct excitatory neuron cell types follows a temporal pattern in which early-born neurons migrate to deep layers (i.e., layers V and VI), while the later-born neurons migrate and surpass early-born neurons to occupy the upper layers (layers II-IV) ^2^. In *Drosophila*, several transcription factors are sequentially explicitly expressed in neural stem cells to control the specification of daughter neuron fates, while very few such transcription factors have been identified in mammals thus far. Using single-cell RNA sequencing (scRNA-seq), Telley and colleagues found that daughter neurons exhibit the same transcriptional profiles of their respective progenitor radial glia, although these apparently heritable expression patterns fade as neurons mature^3^. However, the temporal expression profiles of neural stem cells and the contribution of these specific temporal expression patterns in determining neuronal fate have yet to be wholly clarified in humans and non-human primates. Over the years, non-human primates (NHP) have been widely used in neuroscience research as mesoscale models of the human brain. Therefore, exploring the similarities and differences between NHP and human cortical neurogenesis could provide valuable insight into unique features during human neocortex development.

In mammals, radial glial cells are found in the ventricular zone (VZ), where they undergo proliferation and differentiation. The neocortex of primates exhibits an extra neurogenesis zone known as the outer subventricular zone (OSVZ), which is not present in rodents. As a result of evolution, the diversity of higher mammal cortical radial glia populations increases. Although ventricular radial glia (vRG) is also found in humans and non-human primates, the vast majority of radial glia in these higher species occupy the outer subventricular zone (OSVZ) and are therefore termed outer radial glia (oRG). Outer radial glial (oRG) cells retain basal processes but lack apical junctions ^4^ and divide in a process known as mitotic somal translocation, which differs from vRG ^5^.

VRG and oRG are both accompanied by the expression of stem cell markers such as *PAX6* and exhibit extensive self-renewal and proliferative capacities ^6^. However, despite functional similarities, they have distinct molecular phenotypes. Previous scRNA-seq analyses have identified several molecular markers, including *HOPX* for oRGs, *CRYAB*, and *FBXO32* for vRGs^7^. Furthermore, oRGs are derived from vRGs, and vRGs exhibit obvious differences in numerous cell-extrinsic mechanisms, including activation of the *FGF-MAPK* cascade, *SHH*, *PTEN/AKT*, and *PDGF* pathways, and oxygen (O2) levels. These pathways and factors involve three broad cellular processes: vRG maintenance, spindle orientation, and cell adhesion/extracellular matrix production^8^. Some transcription factors have been shown to participate in vRG generation, such as INSM and TRNP1. Moreover, the cell-intrinsic patterns of transcriptional regulation responsible for generating oRGs have not been characterized.

ScRNA-seq is a powerful tool for investigating developmental trajectories, defining cellular heterogeneity, and identifying novel cell subgroups^9^. Several groups have sampled prenatal mouse neocortex tissue for scRNA-seq ^10,11^, as well as discrete, discontinuous prenatal developmental stages in human and non-human primates, ^7,12^ ^13,14^. The diversity and features of primate cortical progenitors have been explored ^4,6,7,15^.

The temporally divergent regulatory mechanisms that govern cortical neuronal diversification at the early postmitotic stage have also been focused on ^16^. Studies spanning the full embryonic neurogenic stage in the neocortex of humans and other primates are still lacking. Rhesus macaque and humans share multiple aspects of neurogenesis, and more importantly, the rhesus monkey and human brains share more similar gene expression patterns than the brains of mice and humans^17–19^. To establish a comprehensive, global picture of the neurogenic processes in the rhesus macaque neocortex, which can be informative of neocortex evolution in humans, we sampled neocortical tissue at five developmental stages (E40, E50, E70, E80, and E90) in rhesus macaque embryos, spanning the full neurogenesis period. Through strict quality control, cell type annotation, and lineage trajectory inference, we identified two broad transcriptomic programs responsible for the differentiation of deep-layer and upper-layer neurons. We also defined the temporal expression patterns of neural stem cells, including oRGs, vRGs, and IPs, and identified novel transcription factors involved in oRG generation. These findings can substantially enhance our understanding of neocortical development and evolution in primates.

## Results

### scRNA-Seq Analysis of Cell Types in the Developing Macaque Neocortex

In order to establish a comprehensive view of the cellular composition of the rhesus macaque brain at different stages in prenatal period development, we conducted single-cell RNA sequencing (scRNA-seq) on the dissected parietal lobes of eight total rhesus macaque embryos, spanning five developmental stages of prenatal neurogenesis, including E40 (stage of peak neurogenesis) and E50 (Layer 6 formation); as well as at E70 (Layer 5 formation), E80 (Layer 4 formation), and E90 (Layer2-3 formation) (**Figure 1A, Figure S1A**)^2^. We obtained a transcriptome from 53,295 cells after filtering out low-quality cells and removing potential doublets. Each embedding was visualized using uniform manifold approximation and projection (UMAP) of dimension reduction using Seurat, which identified 28 distinct clusters. All cell clusters present in the samples were then annotated (**Figure 1B**), and the respective cell types were identified (**Figure 1C**) based on their expression of molecular markers (**Figure 1D** and **Figure S2, B to C**). According to the expression of the marker genes, we assigned clusters to cell type identities of neurocytes (including radial glia (RG), outer radial glia (oRG), intermediate progenitor cells (IPCs), ventral precursor cells (VP), excitatory neurons (EN), inhibitory neurons (IN), oligodendrocyte progenitor cells (OPC), oligodendrocytes, astrocytes, ventral LGE-derived interneuron precursors and Cajal-Retzius cells, or non-neuronal cell types (including microglia, endothelial, meninge/VALC(vascular cell)/pericyte, and blood cells). Based on the expression of the marker gene, cluster 23 was identified as thalamic cells, which are small numbers of non-cortical cells captured in the sample collection at earlier time points. Each cell cluster was composed of multiple embryo samples, and the samples from similar stages generally harbored similar distributions of cell types.

**Figure 1.**
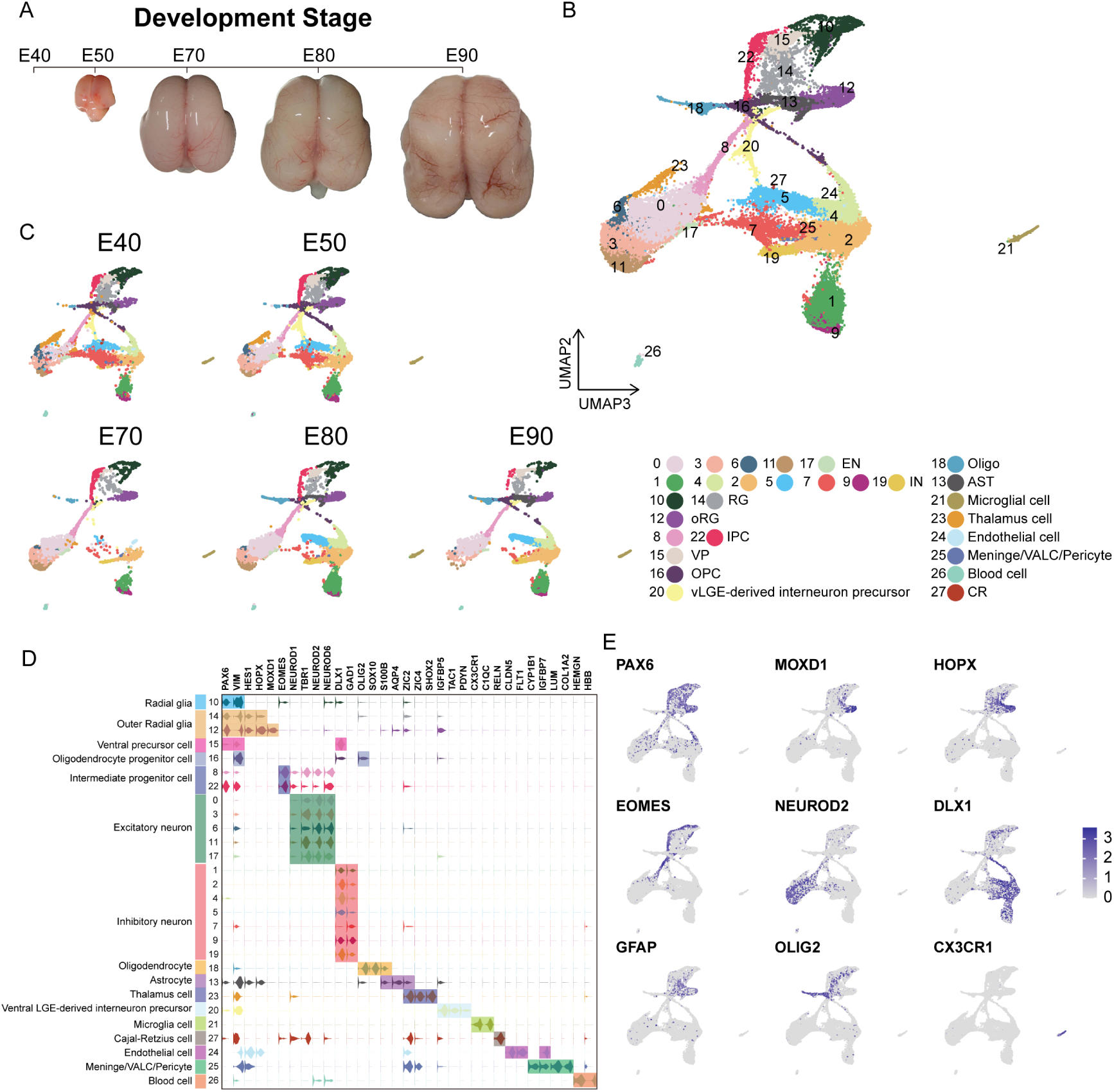
Cell types in macaque prenatal and fetal brain development. (A) Schematic diagram of sample collecting and data analysis. We collected the parietal lobe from the embryos across developmental stages from E40 to E90. (B) and (C) The transcriptome data of single cells were collected and used to do clustering using Seurat. Visualization of major types of cells using UMAP. Dots, individual cells; color, clusters. (D) Violin plot of molecular markers for annotating cell types. (E) The expressions of the classic marker genes for each cell type were plotted for UMAP visualization. Light grey, no expression; Dark blue, relative expression.

In general, ventricular radial glia (vRG) (cluster 10) showed characteristic *VIM* and *PAX6* expression; outer radial glia (oRG) (cluster 12 and cluster 14) highly expressed *HOPX*, previously verified marker^7^; two clusters (cluster 8 and cluster 22) of intermediate progenitor cells (IPCs) that strongly expressed *EOMES*, and one cluster of IPCs (cluster 22) were all topologically close to radial glia, while the other IPC cluster (cluster 8) was closer to neurons, indicating the presence of two stages of IPCs^20^; excitatory neurons were identified by the expression of well-established markers, such as *NEUROD2* and *NEUROD6*; and the astrocyte and oligodendrocyte lineages were identified by *AQP4* and *SOX10* expression, respectively^21^. In addition, we identified *DLX1* and *GAD*-positive cells, which suggested the presence of inhibitory neurons (**Figure 1C** and **Figure S2**).

Collectively, these results suggested that cortical neural progenitors undergo neurogenesis processes during the early stages of macaque prenatal cortical development, while gliogenic differentiation, including oligodendrocytes and astrocytes, occurs in later stages (Figure 1A and 1B).

### Distinct Excitatory Neuronal Types Sequentially Emerge in Developing Cortex

To better understand the temporal dynamics of excitatory neuron (EN) development and differentiation, we compiled a subset of all ENs and re-clustered them into ten subclusters (EN1-10) (**Figure 2A**). We then calculated the relative expression levels of cellular markers for each group and annotated the EN subclusters based on published descriptions of marker function (**Figure S3B**). We found that all EN subclusters could be well-distinguished by differential expression of either deep-layer neuron markers (*BCL11B*, *FEZF2*, and *SOX5*)^22^ or upper-layer neuron markers (*CUX1* and *SATB2*)^23,24^(**Figure 2, B to C**). In previous seminal studies, 3H-thymidine (3H-dT) tracing in macaque rhesus unraveled the sequential generation of cortical neurons, with deep-layer neurons appearing prior to upper-layer neurons^2^. In agreement with this early work, we found that early-born neurons (E40 and E50) predominantly outnumbered later-born neurons in the deep-layer neuron subclusters (EN5 and 10), while upper-layer neuron subclusters contained the inverse proportions (**Figure 2, C and D**).

**Figure 2.**
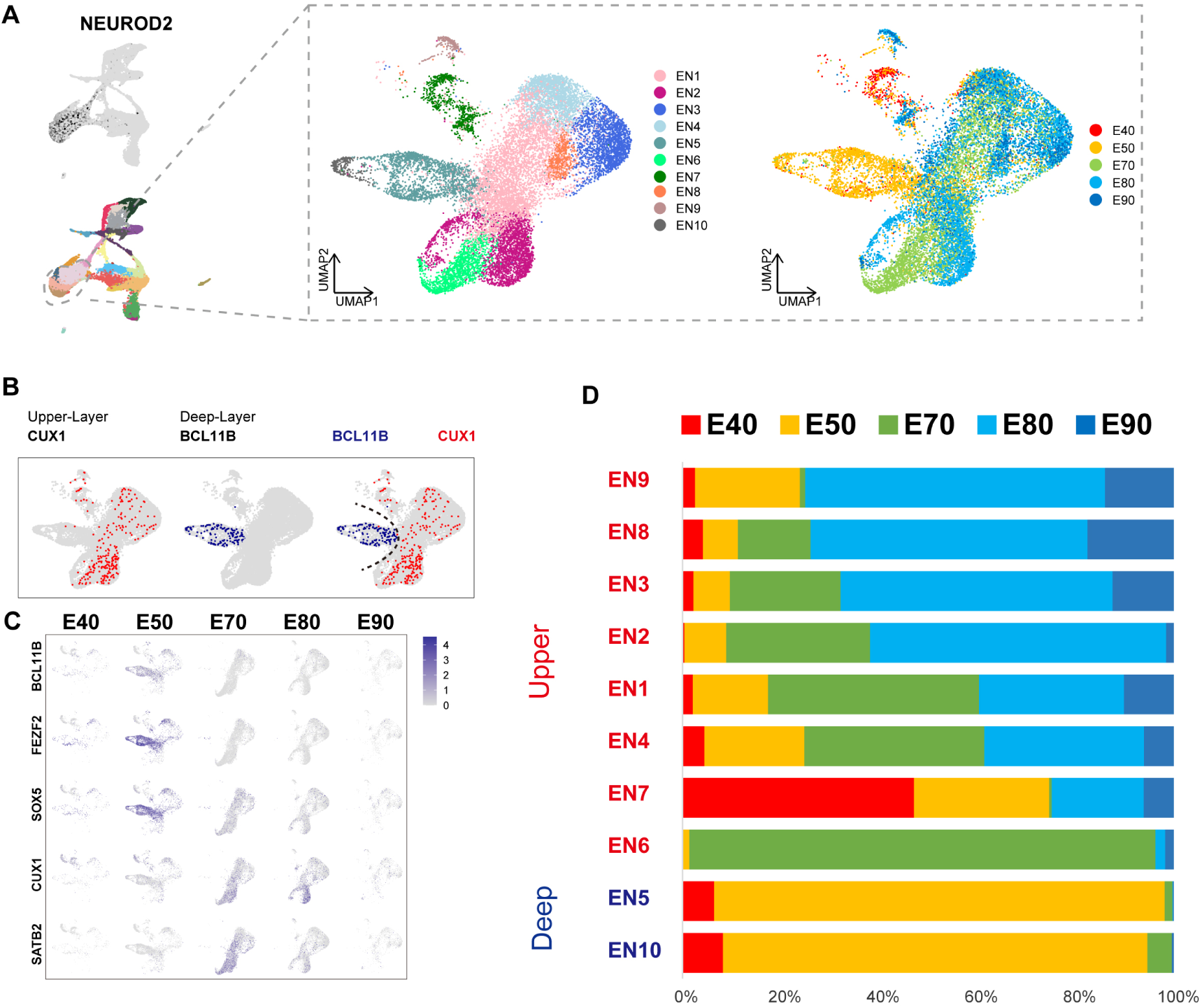
Excitatory neuron subclusters in the developing macaque cerebral cortex. (A) Left, Clustering of excitatory neuron subclusters collected at all time points, visualized via UMAP. Cells are colored according to subcluster identities (left) and collection time points (right). (B) Differentially, the expression of deep-layer marker BCL11B and upper-layer marker CUX1 are highlighted. (C) Excitatory neuron subclusters UMAP plot shows the expression of classic markers for deep layers (*BCL11B*, *FEZF2*, *SOX5*) and upper layers (*CUX1*, *SATB2*) present at each time point. (D) The proportion of different excitatory neuron subclusters corresponding to excitatory neurons in each time point.

We then characterized temporal changes in the composition of each EN subcluster. While the EN 5 and EN 10 (deep-layer neurons) subclusters emerged at E40 and E50 and disappeared in later stages, EN subclusters 1, 2, 3, and 4 gradually increased in population size from E50 to E80, EN subclusters 8 and 9 gradually increased in population size from E80 to E90 (**Figure 2D**). Notably, EN 6 was exclusively found in the cortex at E70 (**Figure S3A**). EN7 is identified as *CUX1*-positive, *PBX3*-positive, and *ZFHX3*-positive excitatory neuron subcluster.

### Specification of Different Progenitor Fates Controlled by Regulator Genes

To focus on the differentiation of neural progenitors, we generated subsets of cell clusters 8, 10, 12, 14, 15, 16, and 22, then annotated them as ventricular radial glia (RG_C10), outer radial glia (oRG_C12 and oRG_C14), intermediate progenitor cells (IPC_C22 and IPC_C8), ventral precursor cells (VP_C15), or oligodendrocyte precursor cells (OPC_C16) (**Figure 3A**). Subclustering analysis revealed that these neural progenitors differentiated into diverse cell types with distinct lineages across the macaque prenatal neocortex (**Figure 3B**). We next investigated trajectories within the neural stem cell pool prior to branching as upper-layer or deep-layer excitatory neurons (EN). The oRG and IPC precursor cell groups exhibited characteristically high specific expression of *HOPX* and *EOMES* respectively.

**Figure 3.**
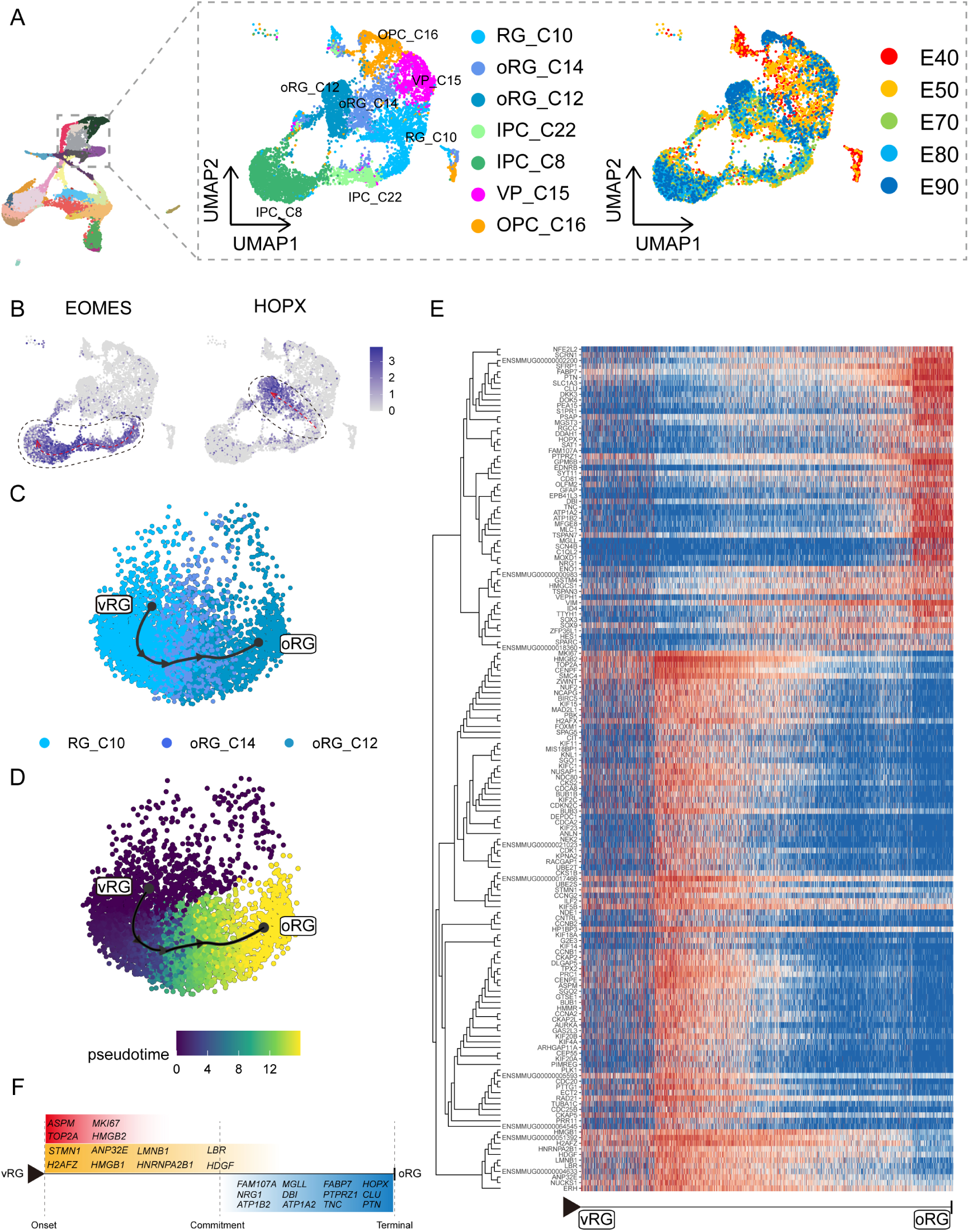
Cell diversity and regulation of progenitor cells in the macaque cortical neurogenesis. (A) UMAP shows eight progenitor clusters and cell annotation. Left cells are colored according to Seurat clusters; right, cells are colored according to the collection time point. (B) Feature plot of outer radial glia marker genes HOPX shows higher expression in C10-C14-C12 (left). Feature plots of Intermediate progenitor cell marker gene EOMES show higher expression in C10-C22-C8 (right). (C) and (D) Pseudotime analysis by Slingshot of HOPX-positive cells (C10-C14-C12). The Slingshot result with the lines indicating the trajectories of lineages and the arrows indicating directions of the pseudotime. Cells are colored according to cell (C) and pesudotime (D). Dots: single cells; colors: cluster and subcluster identity. (E) The heatmap shows the relative expression of top150 genes displaying significant changes along the pseudotime axis of RG to oRG (C10-C14-C12). The columns represent the cells being ordered along the pseudotime axis. (F) Schema diagram of some significant genes related to E. (The depth of the color indicates the levels of gene expression)

Although the presence of oRGs is a well-established feature of primate neurogenesis, and their molecular markers are widely used, the genetic basis and molecular processes leading to their emergence are still poorly understood. To construct gene expression profiles that illustrate the progression from RG_C10 cells to oRGs, we specifically examined changes in expression among RG_C10, oRG_C12, and oRG_C14 cells, then calculated pseudotime trajectories (**Figure 3, C to D**). Based on these trajectories, we then profiled the temporal shifts in the expression of each gene and selected the 150 most significant genes (**Figure 3E**). We found a set of genes that were previously reported as highly expressed in outer radial glia that also showed high expression in the oRG_C12 differentiated pseudotime terminal ^7,25,26^ (such as *SFRP1*, *HOPX*, *FAM107A*, *TNC*, *PTN*, and *MOXD1*), while high *ASPM* expression was typical of cells in the earlier, RG, pseudotime terminal (**Figure 3F**). In addition to these markers, we also found enrichment for some potential regulatory genes at the oRG_C12 terminal, such as Regulator of the cell cycle (*RGCC*), which controls mitotic spindle orientation ^27^, and *TTYH1*, which regulates cell adhesion ^28^. We also found genes that regulate ion channel expression, such as *ATP1A2*, *ATP1B2*, and *SCN4B*, which were not previously known to participate in oRG differentiation.

Analysis of DEGs specific to the RG pseudotime terminal identified cell cycle-related genes such as *MKI67*, *TOP2A*, *CDC20*, and *CCNA2*. Within the glia-like cells that largely comprised the oRG_C12 terminal, we also detected a population of true glial genes that exhibited high expression of *SLC1A3*, *ZFP36L1* ^29^, *GFAP*, *DBI*, and *EDNRB* ^30^. In addition, our analyses uncovered several DEGs that have not yet been investigated for a role in radial glia function and differentiation, such as *DKK3*, *DDAH1*, *SAT1*, and *PEA15*. Finally, we screened for significant DEGs associated with RG-to-IPC and OPC-to-astrocyte/oligodendrocyte differentiation trajectories (Figures **S5 and S6**).

### The Generation of Deep-layer and Upper-layer Neurons Follows Distinct Terminal Trajectories

Based on our finding above that the deep-layer neurons appear earlier than upper-layer neurons, we next sought to characterize dynamic shifts in the expression of transcriptional regulators that potentially contribute to determining neuronal fates. To this end, we performed pseudotime trajectory analysis of EN lineages (including deep-layer and upper-layer neurons) (**Figure 4A**), then superimposed the cluster labels from RG_C10, oRG_C12, and oRG_C14 stem cells in UMAP plots (Figure 3A) and early (E40 and E50) and late (E70, 80 and 90) emergence stages (Figure 2A) over the extracted transcriptomic data. We then calculated the RG-specific differentially expressed genes (rgDEGs) and the DEGs of early and late neuron-specific (nDEGs). The set of genes that overlapped between rgDEGs and nDEGs (termed mapping genes) can thus be used to map neuronal subtypes to different neural stem cell populations (**Figure. S4**). We hypothesized that mapping genes, such as *FEZF2* and *DOK5*, may function in radial glial cells to specify neuronal progeny (**Figure 4B**). Our results provide single-cell transcriptome data to support that apical progenitors and daughter excitatory neurons share molecular temporal identities during neocortex prenatal corticogenesis ^31–33^.

**Figure 4.**
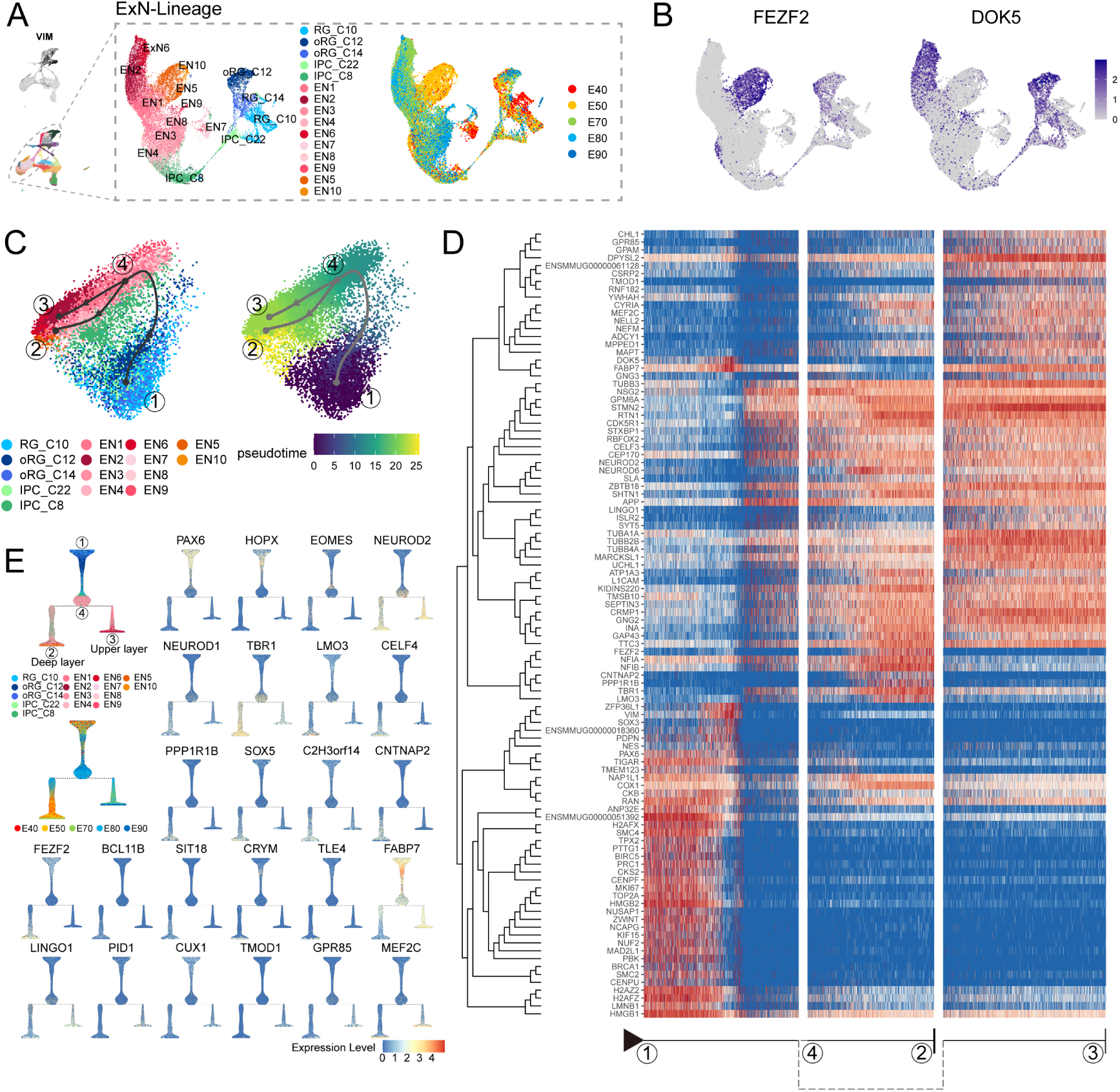
Transcriptional regulation of excitatory neuron lineage during prenatal cortical neurogenesis. (A) UMAP shows the alignment of macaque cortical NPCs, IPCs, and excitatory neurons. Left, cells are colored according to cell annotation. Different yellow/orange colors are used for deep-layer excitatory neuron subclusters (EN5 and EN10), and different red/pink colors are used for upper-layer excitatory neuron subclusters (EN1, EN2, EN3, EN4, EN6, EN7, EN9 and EN10). Right, cells are colored according to the time point of collection. (B) Dot plot showing the marker genes for the deep-layer excitatory neuron (*FEZF2*) and upper-layer excitatory neuron (*DOK5*). Light grey, no expression; Dark blue, relative expression. (C) Pseudotime analysis by Slingshot projected on PCA plot of RGCs, oRGCs, IPCs, and excitatory neuron subclusters. The Slingshot result indicates the trajectories of lineages, and the arrows indicate the directions of the pseudotime. Dots: single cell; colors: cluster and subcluster identity. Framed numbers marked the start point, endpoint, and essential nodes of the Slingshot inference trajectory. Framed number “1” was the excitatory neuron lineage trajectory start point (C10). Framed number “4” marked immature neurons. Framed numbers “2” and “3” marked deep-layer and upper-layer neurons. Cells are colored according to cell annotation and pseudotime. (D) The heatmap shows the relative expression of the top 100 genes displaying significant changes along the pseudotime axis of each lineage branch. The columns represent the cells being ordered along the pseudotime axis. (E) Left, Slingshot branching tree related to Slingshot pseudotime analysis in C. The root is E40 earliest RG (C10), tips are deep-layer excitatory neurons generated at the early stage (E40, E50), and upper-layer excitatory neurons are generated at the later stage (E70, E80, E90). Right, branching trees showing the expression of marker genes of apical progenitors (*PAX6*), outer radial glia cells (*HOPX*), intermediate progenitors (*EOMES*), and excitatory neurons (*NEUROD2*), as well as genes in Figure 3E, including callosal neurons (*SATB2*, *CUX2*), deeper layer neurons (*SOX5*, *FEZF2*), corticofugal neurons (*FEZF2*, *TLE4*). There is a sequential progression of radial glia cells, intermediate progenitors, and excitatory neurons.

Since substantial evidence indicates that neuronal fate is determined post-mitotically in the mammalian neocortex, analysis of differentially expressed genes throughout the differentiation process could reveal the genetic mechanisms responsible for deciding stem cell fate as different neuronal progeny. Using Dynverse R packages, we constructed pseudotime trajectories for the cluster data, which identified a bifurcating trajectory from neural stem cells (including RG_C10, oRG_C12, and oRG_C14) that leads to either deep-layer neurons in one branch or to upper-layer neurons in the alternative branch (**Figure 4C**). However, it should be noted that the two distinct fates share a common path from neural stem cells to immature neurons (Figure 4C, from **dot1** to **dot4**), supporting the likelihood that neuronal fate is primarily determined post-mitotically.

Based on this trajectory, we categorized the DEGs that exhibit dynamic, temporal shifts in expression over pseudotime, resulting in broad clusters. The first of these clusters was enriched with neural stem cells, characterized by high expression of *PAX6*, *VIM*, and *TOP2A*, which likely participate in regulating stemness and proliferation. The three remaining DEG clusters were enriched in either deep-layer neurons, upper-layer neurons, or both (**Figure 4D**). More specifically, we found that most DEGs, such as *STMN2*, *TUBB3*, *NEUROD2*, and *NEUROD6*, were shared between both branches, illustrating the differentiation processes between deep and upper-layer ENs. We also identified deep layer-specific DEGs (including *FEZF2*, *TBR1*, and *LMO3*) and upper layer-specific genes (including *MEF2C* and *DOK5*). We then organized the cell types into a branching tree based on their differential expression of these marker genes, including stem-cell genes *PAX6*, *HOPX*, and *EOMES*, in addition to the above deep-layer-and upper-layer-specific genes (**Figure 4E**). The separation into distinct, branch-specific sets of DEGs suggested a role for these cell type-specific transcription factors and other genes in terminal neuron differentiation.

### Conserved and Divergent Features of Human, Macaque and Mouse Neocortical Progenitor Cells

Next, we performed liger integration ^34^ to integrate the macaque single-cell dataset with published mouse ^11^ and human datasets, which were created by combing two published human scRNA-seq data of 7 to 21 postconceptional week (PCW) neocortex ^35,36^, to conduct a cross-species comparison. Species-integrated UMAP showed that all major cell types were well integrated between the three species (**Figure 5A**).

**Figure 5.**
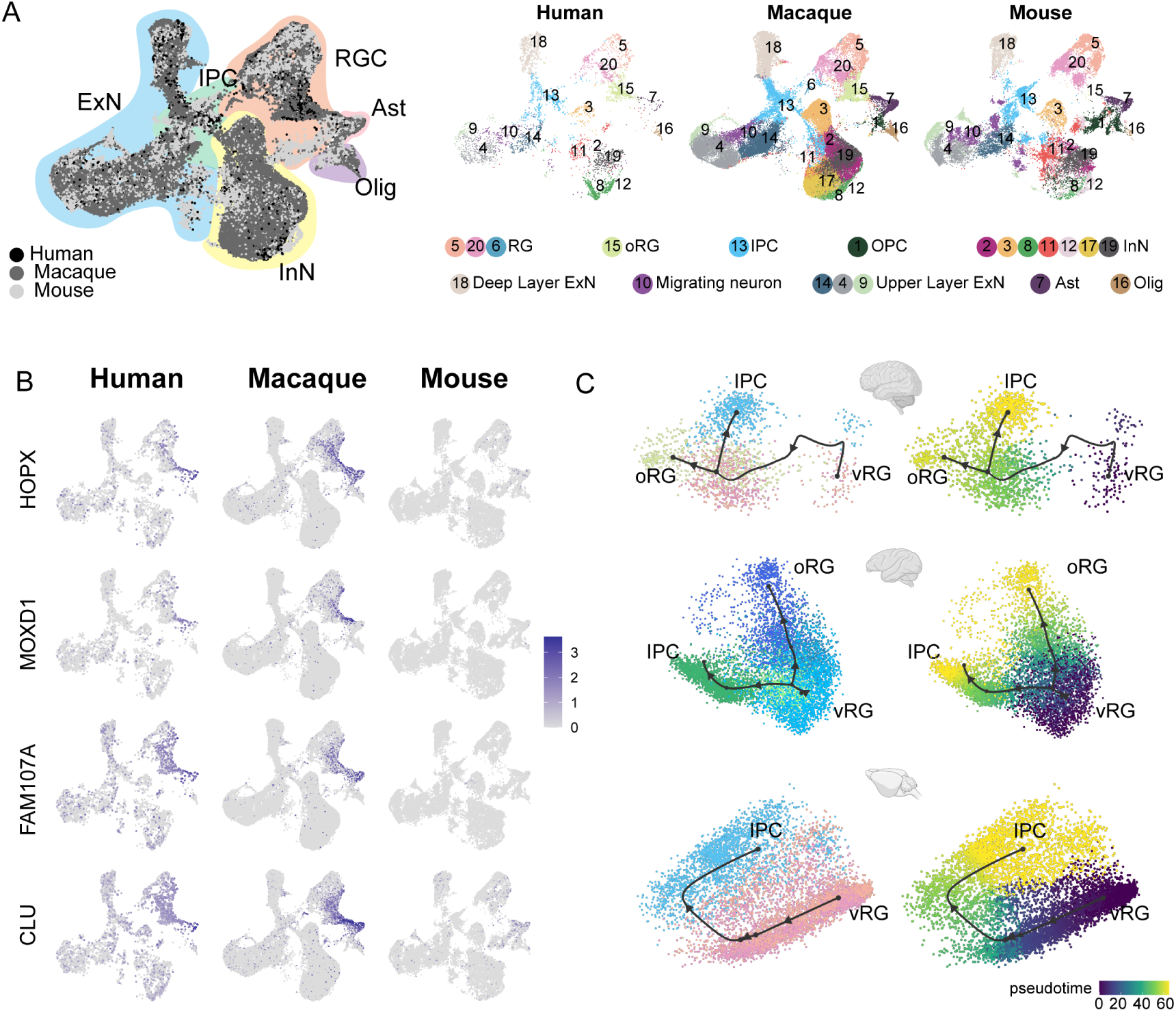
Integration of human, macaque and mouse single-cell datasets reveals conserved and divergent progenitor cell types. (A) Left: UMAP plot of cross-species integrated single-cell transcriptome data with liger. Colors represent different major cell types (Black: Human dataset; Dark grey: macaque dataset; Liger grey: mouse). Right: The UMAP plot of each dataset, colored by liger cluster. (B) The expressions of the classic oRG marker genes were plotted to UMAP visualization. Light grey, no expression; Dark blue, relative expression. (C) Comparison of vRG→oRG and vRG→ IPC developmental trajectories between human, macaque, and mouse.

It is clearly observed that similar cellular processes with comparable temporal progression sequentially generate deep-layer and upper-layer neurons in both primates and rodents (**Figure S7, A to C**). However, in primates, the number and relative development period of upper neurons are more extended than that of rodents (**Figure S7D**). Besides, another important evolutionarily divergent feature in cortical development between primates and rodents is the abundance of oRG cells in primates’ outer subventricular zone (oSVZ) (human and macaque). Our human-macaque-mouse cross-species analysis showed that strong expression of the oRG marker genes, such as *HOPX*, *MOXD1*, *FAM107A* and *CLU* (3) in both human and macaque datasets (**Figure 5B**), while barely detectable expression in mouse dataset, which is consistent with previous reports ^37^.

Furthermore, We picked *HOPX*-positive and *EOMES*-positive cells (ligercluster 5, 20, 13, 15) from the human, macaque, and mouse datasets and performed developmental trajectories analysis with slingshot (**Figure 5C**). Comparing vRG to IPC trajectory between human, macaque, and mouse, we found this biological process of vRG-to-IPC is remarkably conserved across species. However, the vRG to oRG trajectory is divergent between species because the oRG population was not identified in the mouse dataset. The latter process is almost invisible in mice but similar in humans and macaques.

Previous studies have shown that neural progenitors exhibit distinct temporal expression patterns during neurogenesis in mice. However, similar temporal profiling has yet to be thoroughly investigated in macaque neural stem cells. Based on its topological position and associated markers that suggest it functions as a developmental root, we examined the temporal expression profile of RG_C10 cells (Figure 3A). We first identified marker genes to distinguish RG_C10 cells in different prenatal stages (i.e., E40-E90) and used these markers to generate dynamic expression profiles. We then clustered these dynamically expressed macaque genes into five types (**Figure 6A**), consisting of Type 1 and 2 genes that decreased in expression during neurogenesis, Type 3, 4 and 5 genes that gradually ramped up in expression throughout neurogenesis. We then performed a similar profiling of dynamic gene expression using data obtained from mouse apical progenitors^10^ and categorized the DEGs into the same five categories. A total of 72 homologous transcription factors were shared between human, macaque and mouse transcriptomes. Analysis of their temporal dynamics revealed that they exhibited similar temporal patterns between species such as *NEUROD1*, *NEUROG2*, *POU3F2*, *ETV1*, *ETV5* and *FOS* (**Figure 6B** and **Figure S8**), which indicated that the majority of transcription factors had conserved temporal dynamics between species, and thus evolutionarily conserved patterns of regulation in neural stem cell differentiation.

**Figure 6.**
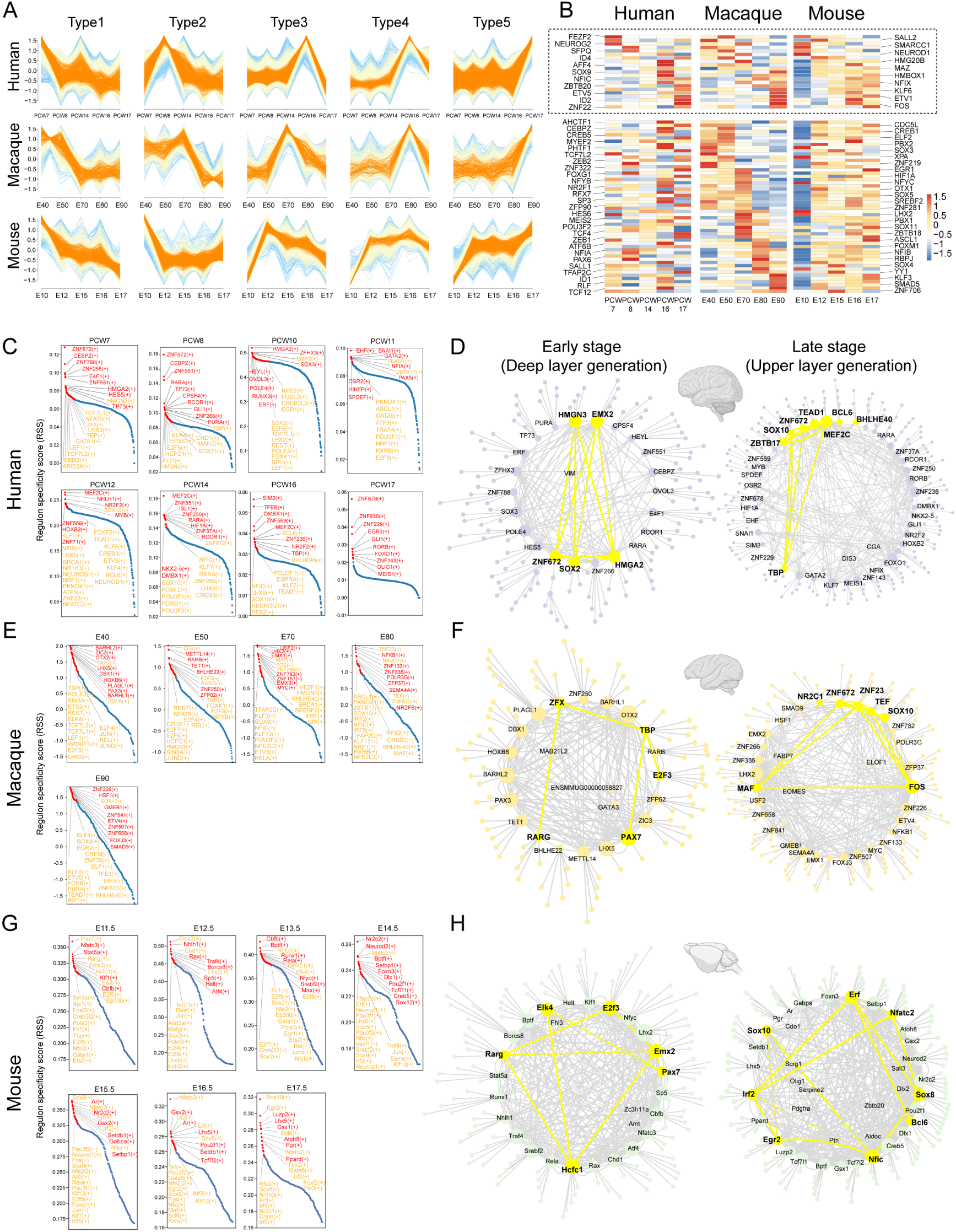
The patterns of transcriptional regulation comparative analysis responsible in vRGs. (A) Normalized expressions of genes that show temporal dynamics in the vRGs of human, macaque and mouse. (B) Temporal expression heatmap of homologous transcription factor genes among human, macaque, and mouse temporal expression heatmap. (The TFs genes in the dashed boxes showed similar temporal expression patterns across species. (C), (E) and (G) Regulon specificity score for each timepoint in human, macaque, and mouse vRG. Regulons with high scores in multiple species vRG cells are colored yellow. (D), (F) and (H) show a network generated with Cytoscape using the top 10 regulons in the human, macaque, and mouse vRG at each time point and their top 5 target genes identified by SCENIC as an input. The interactions between conserved TFs in more than one species are colored yellow.

To identify the master regulators (MRs) related to cortical neurogenesis among macaque and mouse, we used the SCENIC workflow to analyze the gene regulatory networks of each transcription factor (**Figure 6, C** to **H**). Among all the regulatory pairs associated with each prenatal stage, we selected ten TFs with the highest regulon specificity scores (**Figure 6 C, E and G**) and their top 5 target genes, as visualized by Cytoscape. Analysis of the regulatory activity of human, macaque and mouse^11^ prenatal neocortical neurogenesis indicated that despite commonalities in the roles of classical developmental TFs such as *GATA1*, *SOX2*, *HMGN3*, *TCF7L1*, *ZFX*, *EMX2*, *SOX10*, *NEUROG1*, *NEUROD1* and *POU3F1*. The top 10 TFs of the human, macaque, and mouse vRG each time point and their top 5 target genes identified by pySCENIC as an input to construct the transcriptional regulation network (**Figure 6 D, F** and **H**). Some conserved regulatory TFs present in more than one species are identified, such as *HMGN3*, *EMX2*, *SOX2*, and *HMGA2* at an early stage when deep-lager generation and *SOX10*, *ZNF672*, and *ZNF672* at a late stage when upper-lay generation.

## Discussion

The cerebral cortex region of the brain is responsible for extraordinary cognitive capacities, such as abstract thinking and language. The neocortex of non-human primate rhesus macaque resembles that of humans in many aspects. Thus, a comprehensive investigation of macaque neurogenesis can improve our understanding of neocortical development and evolution. For this purpose, we performed scRNA-seq in prenatal neocortical tissues of macaque. Our data support previous omics studies of prenatal macaque brain development ^38^ ^39^ ^17^ and recently published human neocortex development studies ^40^. We constructed a single-cell resolution transcriptomic atlas of the developing macaque neocortex, which we then used to identify dynamically expressed genes that likely contribute to the maturation of distinct neuron types, temporal expression patterns of neural stem cells, and the generation of oRGs.

The existence of oRGs has long been demonstrated in humans and primates through live imaging and immunostaining ^41^. ORGs share similarities in their patterns of gene expression with vRGs (e.g., *SOX2*, *PAX6*, and *NESTIN*) but also specifically express several genes (such as *MOXD1*, *HOPX*, and *FAM107A*) ^7,42^. Recently, the molecular mechanisms associated with the generation and amplification of oRGs were shown to include signaling pathways such as the *FGF-MAPK* cascade, *SHH*, *PTEN/AKT*, and *PDGF* pathways, as well as proteins such as *INSM*, *GPSM2*, *ASPM*, *TRNP1*, *ARHGAP11B*, *PAX6*, and *HIF1α*. A number of these proteins were validated by genetic manipulation of their activities or expression in mouse, ferret, and marmoset ^43–46^. The dynamically expressed genes and pseudotime trajectories from vRGs to oRGs presented in this work are in agreement with these previous reports. In addition to known regulators, we detected differential expression of ion channel regulators in oRGs, such as *ATP1A2*, *ATP1B2*, and *SCN4B*, suggesting that hyperpolarization and depolarization processes may participate in promoting or maintaining oRG populations. Indeed, hyperpolarization in mouse apical progenitors has been shown to promote the generation of IPCs and indirect neurogenesis ^47^, although it remains unclear if similar mechanisms contribute to the development of the primate neocortex.

The sequential generation of cortical neurons has long been observed in both rodents and primates, and many transcription factors are known to drive post-mitotic specification of neuronal progeny, mainly in rodents. Here, we found genes conserved across species, such as *FEZF2* and *TBR1*, involved in specifying deep-layer neurons in the macaque neocortex. The slingshot pseudotime analysis, which reflected the expression pattern of genes during appropriate lineage trajectories, suggests that some genes are lineage-restricted or even involved in lineage fate determination. For example, *FEZF2*, *PPP1R1B* (also known as *DARPP32*), *SOX5*, *LMO3*, and *CELF4* may be involved in the fate specialization of deep neurons. In contrast, *TMOD1*, *PID1*, *LINGO1*, *CRYM*, and *MEF2C* for upper-layer neurons. Notably, *FEZF2* ^48^ and *SOX5* ^49^ have been confirmed associated with deep layer neuron specification in mice, while others, such as *TMOD1*, *PID1*, and *LINGO1*, have not been reported before, and their potential functions in the fate specification of cortical projection neurons need to be further explored. To date, only a very limited set of genes have been shown to function in specifying neuronal progeny in progenitor cells, which suggests that regulation in progenitors is more complex than in neurons, where a single transcription factor can trigger a transcriptional cascade to promote the maturation of neurons. In the current study, we found two types of genes (Dynamic type 1 and 5), which have gradually altered expression patterns along with the neurogenic stage, indicating a transition in cell state in both vRGs and oRGs. In addition to these gradual transition genes, we also identified sets of genes (Dynamic type 2-4) with sharp changes in expression during neurogenesis, which likely function in specifying different neuronal subtypes. These findings led us to postulate that a combination of biological processes and pathways change over time to coordinate gradual transitions from neural stem cells to daughter neurons.

This study also faces limitations. During the macaque prenatal neurogenesis, dorsal radial glia cells generate excitatory neurons by direct and indirect neurogenesis. In general, the number of progenitor cells gradually decreases throughout development, with progenitor cells generating deep neurons in the early stage of development and upper excitatory neurons in the later stages. We observed the disappearance of two subclusters of ENs (EN 5 and EN 10) from later-stage samples (Figure S3). This disappearance could also be explained by the development of axons and dendrites in early-born neurons, which have higher morphological complexity and greater vulnerability. Due to the fixed size of the BD Rhapsody microwells, this sing-cell capture method might be less efficient in capturing mature excitatory neurons but has a good capture effect on newborn neurons at each sampling time point. Nonetheless, this transcriptomic atlas uncovers the molecular signatures of the primary cell types and temporal shifts in gene expression of differentiating progenitor cells during neocortex layer formation, thus providing a global perspective into neocortical neurogenesis in the macaque.

### Materials and Methods Animal

Animals and frozen tissue samples from prenatal rhesus macaque (Macaca mulatta) were provided by the National Medical Primate Center, Institute of Medical Biology, Chinese Academy of Medical Sciences. Timed pregnancy-derived biological replicate specimens were profiled at each prenatal developmental stage (E40, E50, E70, E80 and E90)^50^. These time points were selected to coincide with peak periods of neurogenesis for the different layers of the neocortex based on previous studies ^50,51^. All animal procedures followed international standards and were approved in advance by the Ethics Committee on Laboratory Animals at IMBCAMS (Institute of Medical Biology, Chinese Academy of Medical Sciences).

### Fetal brain sample details

We collected eight pregnancy-derived fetal brains of rhesus macaque (Macaca mulatta) at five prenatal developmental stages (E40, E50, E70, E80, E90) and dissected the parietal lobe cortex. Because of the different development times of rhesus monkeys, prenatal cortex size and morphology are different. To ensure that the anatomical sites of each sample are roughly the same, we use the lateral groove as a reference to collect the parietal lobe for single-cell sequencing (as indicated by bright yellow in Figure S1A) and do not make a clear distinction between the different regional parts including primary somatosensory cortex and association cortices in the process of sampling.

### Cell preparation

We followed the BD Cell Preparation Guide to wash, count, and concentrate cells in preparation for use in BD Rhapsody™ System Whole transcriptome analysis.

### Single-cell RNA-seq data processing

Single-cell capture was achieved by random distribution of a single-cell suspension across >200,000 microwells through a limited dilution approach. Beads with oligonucleotide barcodes were added to saturation so that a bead was paired with a cell in a microwell. The cell-lysis buffer was added so that poly-adenylated RNA molecules hybridized with the beads. Beads were collected into a single tube for reverse transcription. Upon cDNA synthesis, each cDNA molecule was tagged on the 5′ end ( the 3′ end of a mRNA transcript) with a molecular index and cell label indicating its cell of origin. Whole transcriptome libraries were prepared using the BD Resolve single-cell whole-transcriptome amplification workflow. In brief, the second strand of cDNA was synthesized, followed by ligation of the adaptor for universal amplification. Eighteen cycles of PCR were used to amplify the adaptor-ligated cDNA products. Sequencing libraries were prepared using random priming PCR of the whole-transcriptome amplification products to enrich the 3′ ends of the transcripts linked with the cell label and molecular indices. Sequencing was performed with Illumina NovaSeq 6000 according to the manufacturer’s instructions. The filtered reads were aligned to the macaque reference genome file with Bowtie2 (v2.2.9). Macaque reference genome was downloaded from Ensemble database (ftp://ftp.ensembl.org/pub/release100/fasta/macaca_mulatta/dna/Macaca_mulatta.Mmul_10.dna.toplevel.fa.gz, ftp://ftp.ensembl.org/pub/release-100/gtf/macaca_mulatta/Macaca_mulatta.Mmul_10.100.gtf.gz). We analyzed the sequencing data following BD official pipeline and obtained the cell-gene expression matrix file.

### QC and Data analysis

To mitigate the effect of over-estimation of molecules from PCR and sequencing errors, we used the Unique Molecular Identifier (UMI) adjustment algorithms recursive substitution error correction (RSEC) and distribution-based error correction (DBEC), which were contained in the BD Rhapsody™ pipeline. Quality control was applied based on the detected gene number and the percentage of counts originating from mitochondrial RNA, ribosomal RNA, and hemoglobin gene per cell. Then, cells were filtered to retain only higher quality (with less than 7.5% mitochondrial gene counts, less than 5.5% ribosomal gene counts, and detected genes above 400 and less than 6000). Additionally, we identified cell doublets using Scrublet v0.142 with default parameters and removed doublets before data analysis. After quality control, a total of 59,045 cells remained for subsequent analysis.

To integrate cells into a shared space from different datasets for unsupervised clustering, we used SCTransform workflow in R packages Seurat^52^ V4.0.3 to normalize the single-cell RNA-seq data from different samples. We identified the variable features of each donor sequencing data using the “FindVariableFeatures” function. A consensus list of 2000 variable genes was then formed by detecting the most significant recovery rates genes across samples. Mitochondrial and ribosomal genes were not included in the variable gene list. Next, we used the SCTransform workflow in Seurat to normalize the single cell-seq data from different samples. During normalization, we also removed confounding sources of variation, including mitochondrial mapping and ribosomal mapping percentages. Standard RPCA workflow was used to perform the integration.

With a “resolution” of 0.5 upon running “FindClusters,” we distinguished major cell types of nerve cells and non-nerve cells in the umap according to known markers, including radial glia cell, outer radial glia cell, intermediate progenitor cell, excitatory neuron, inhibitory neuron, astrocyte, oligodendrocyte, Cajal-Retzius cell, microglia, endothelial cell, meningeal cells, and blood cells, which were distinguished on the first level. By following a similar pipeline, subclusters were identified. We finalized the resolution parameter on the FindClusters function once the cluster numbers did not increase when we increased the resolution. Then, we checked the differentially expressed genes (DEGs) between each of the clusters using the “FindAllMarkers” and “FindMarkers” functions with logfc.threshold=0.25.

### Construction of single-cell developmental trajectory

Single-cell developmental lineage trajectories construction and discovery of trajectory transitions were performed using Slingshot(V 2.2.0) ^53^, from the Dyno (V 0.1.2) ^54^ platform, with PCA dimensionality reduction plot results. The direction of the developmental trajectory was adjusted by reference to the verified relevant studies.

### Cross-species transcriptome data integration and analysis

Mouse datasets were downloaded from the Gene Expression Omnibus (GEO SuperSeries GSE153164) and at the Single Cell Portal: https://singlecell.broadinstitute.org/single_cell/study/SCP1290/molecular-logic-of-cellular-diversification-in-the-mammalian-cerebral-cortex. Human datasets were download from the Gene Expression Omnibus (GEO) under the accession number GSE104276 and GSE104276. We created a human database by combining the two published human prenatal cortical development datasets of 7 to 21 postconceptional weeks (PCW) into one containing cell numbers comparable to our macaque and published mouse dataset using the single cell transcriptome analysis pipeline of R package Seurat. We used the biomaRt package to convert the gene symbols in the mouse and macaque expression matrices into their human homologs. To perform cross-species analysis, we used the LIGER ^[31]^ method to integrate the macaque single-cell dataset with the mouse and human datasets. We ran the “optimizeALS” function in rliger to perform integrative non-negative matrix factorization(iNMF) on the scaled species-integrated dataset (k = 20, lambda = 5, max.iters = 30). Species-integrated UMAP showed that all major cell types were well integrated between the three species (Figure 5A). We used the “calcAlignment” function in rliger to quantify how well-aligned datasets are and got the metric of 0.891231545756746, which was very close to “1”. This suggested LIGER well-integrated cross-species datasets.

### Transcription Factor Regulatory Network Analysis

We inferred regulon activities for vRG cells of human, macaque and mouse using pySCENIC ^[56]^ workflow (https://pyscenic.readthedocs.io/en/latest/tutorial.html) with default parameters. Since there is no transcriptional regulator database for macaques, macaques’ genome is similar to the human genome. When we analyzed the macaque dataset, we used the human transcriptional regulator database for pySCENIC analysis. GRNboost2 in scenic was used to infer gene regulatory networks based on co-expression patterns. RcisTarget^55^ was used to analyze the TF-motif enrichment and direct targets among different species databases. The regulon activity for each developmental time point was identified using AUCell, and the top enriched activated TFs were ranked by –log10(p_value). Lastly, we evaluated the interaction networks analysis of TFs and their target genes by Cytoscape^56^ (version 3.8.1).

### Mfuzz Clustering

Firstly, we picked up vRG cells in each species dataset and screened the differentially expressed genes (DEGs) between adjacent development time points using the “FindMarkers” function (with min.pct = 0.25, logfc.threshold = 0.25). After separate normalization of the DEG expression matrix from different species datasets, we use the “standardise” function of mfuzz to perform standardization. The DEGs of vRG in each species were grouped into five different clusters using the Mfuzz package in R with fuzzy c-means algorithm^57^.

### Transcription Factor and RNA binding protein analysis

Mouse, macaque, and human transcription factor (TF) gene lists were downloaded from AnimalTFDB (http://bioinfo.life.hust.edu.cn/AnimalTFDB/). Mouse, macaque, and human RNA binding protein (RBP) gene lists were downloaded from EuRBPDB (http://EuRBPDB.syshospital.org). Heatmaps of the TFs and RBPs expression profiles were generated with normalized and standardized DEGs expression matrixes from mfuzz.

## Acknowledgments

Funding: This work was supported by CAMS Innovation Fund for Medical Sciences (CIFMS, 2021-I2M-1-024, 2021-I2M-1-019) and National Science and Technology Innovation 2030 Major Program 2021ZD0200900.

## Data availability

The raw sequence data reported in this paper have been deposited in the Genome Sequence Archive in National Genomics Data Center, China National Center for Bioinformation / Beijing Institute of Genomics, Chinese Academy of Sciences (GSA: CRA007718) that are publicly accessible at https://ngdc.cncb.ac.cn/gsa. Codes used to analyze results in this paper are available on GitHub at https://github.com/cheneylemon/developing-macaque-neocortex.

## Author Contributions

X.P. and B.Q. supervised the research. P.S. and Y.C. conceived of the experimental design and bioinformatics analysis. Y.Z., Z.H. and S.L. established the macaque prenatal developmental model. W.L. performed the anatomical analysis of monkeys and related tests. L.X. and J.Z. contributed to drafting or revising the manuscript and figures. Z.Y. and L.X. performed single-cell transcriptome data analysis.

## Competing Interest Statement

The author(s) declared no potential conflicts of interest with respect to the research, authorship, and/or publication of this article.

## Supporting Information

**Figure S1.**
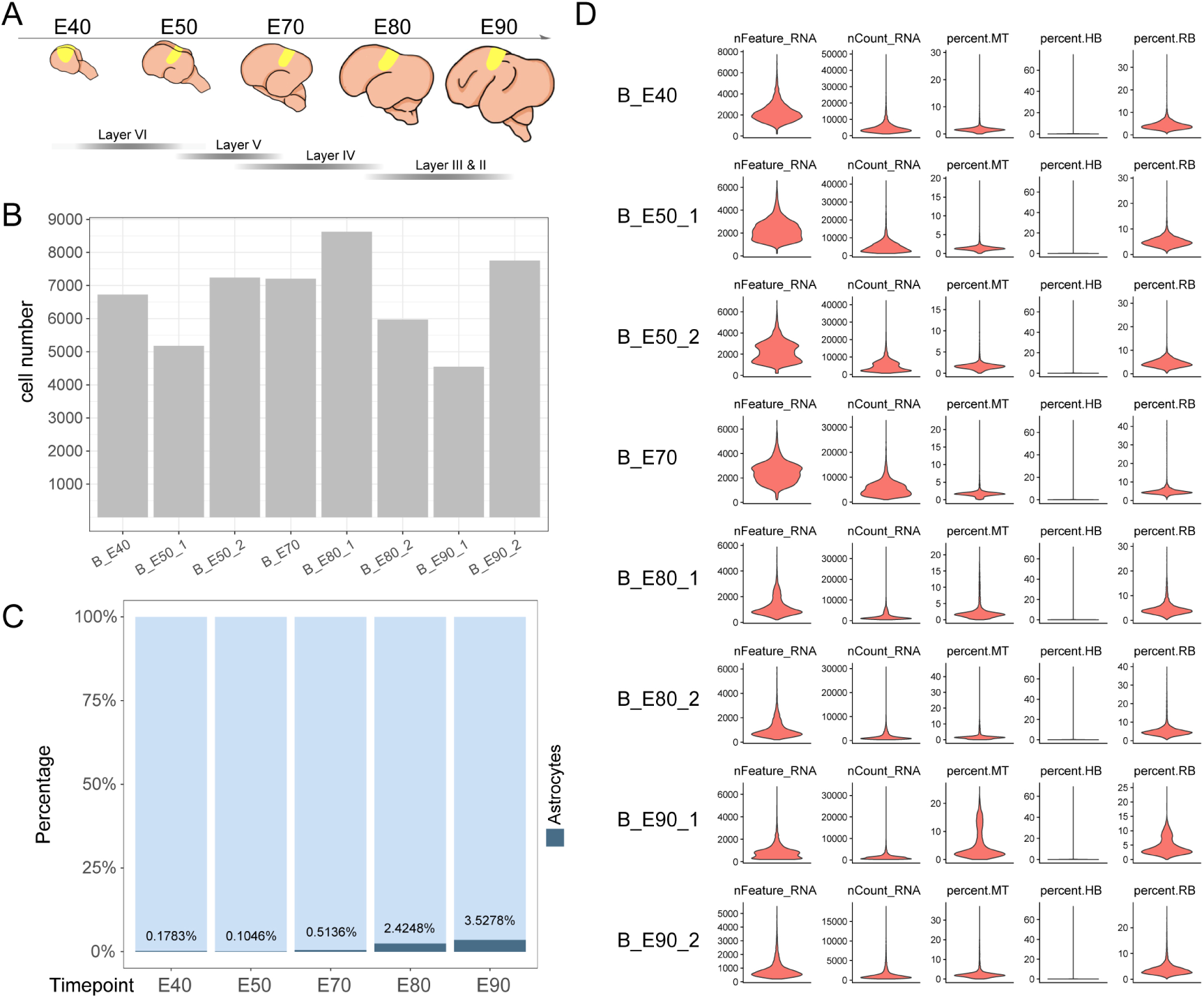
Sample collection and Quality Control. **(A)**Schematic diagram of sample collecting anatomical area. **(B)**The cell number per sample after quality control. **(C)** Bar chart of astrocytes proportion statistical at each time point. **(D)** Single-cell transcriptome library information for each sample.

**Figure S2.**
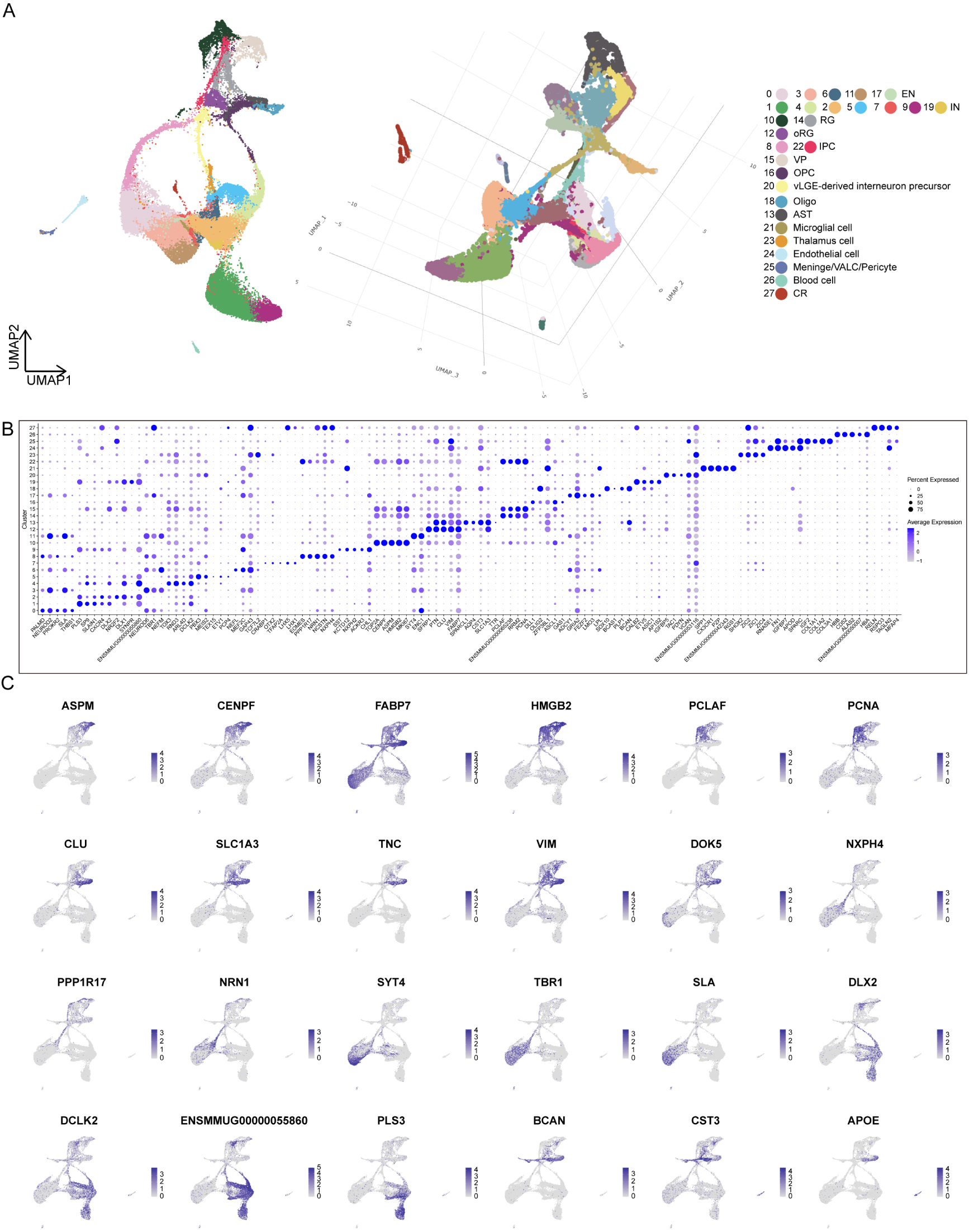
scRNA-Seq uncovers cell type in the developing macaque neocortex. **(A)** Visualization of different dimensionality reduction of all cells. (Left, UMAP visualization with UMAP1 and UMAP2; right, 3D model of UMAP visualization with UMAP1, UMAP2, and UMAP3). **(B)** Top marker genes for each of the 28 cell clusters shown in Figure 1B. **(C)** Feature plots of marker gene expression. Colors represent scaled gene expression.

**Figure S3.**
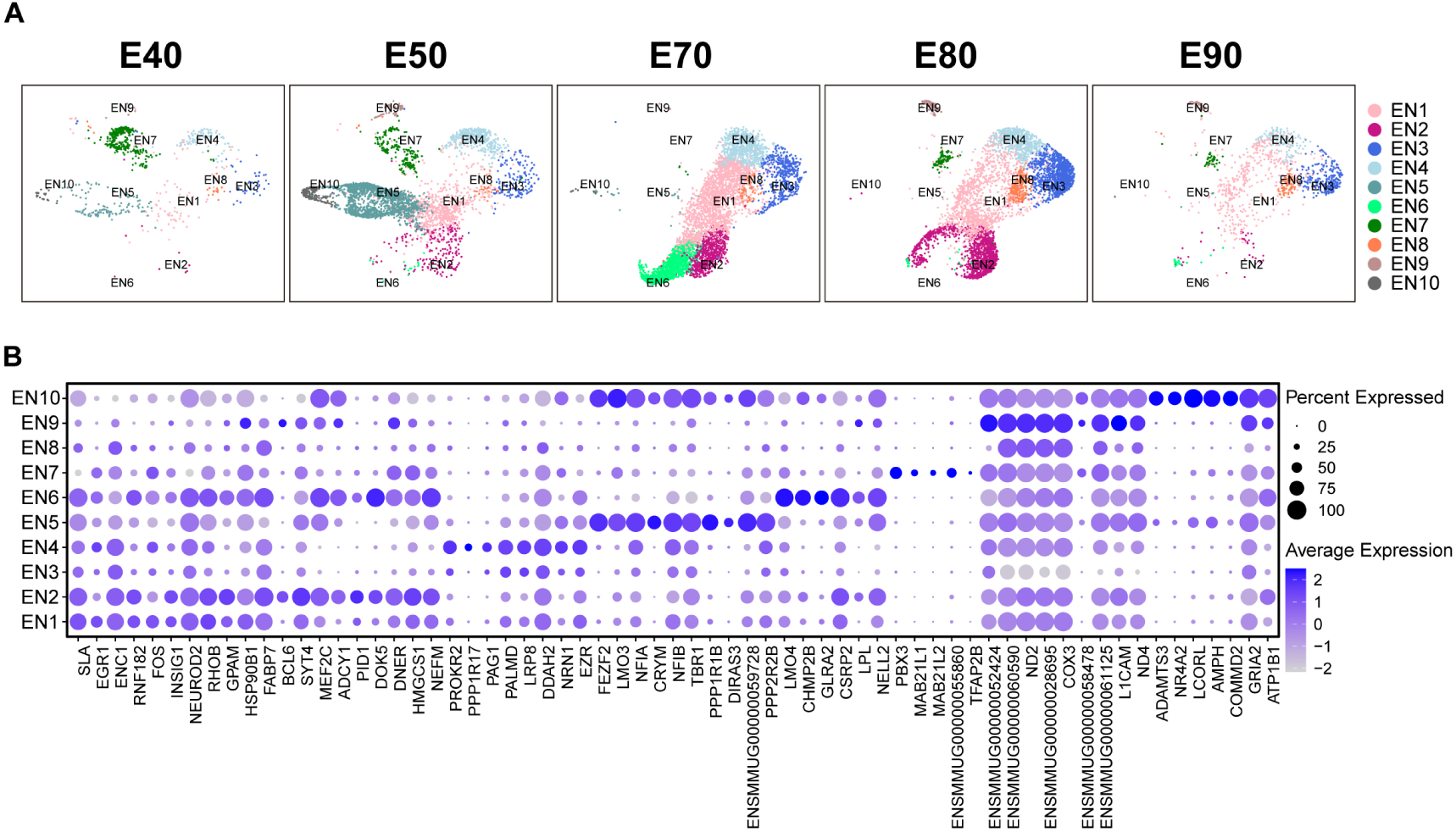
Additional information for excitatory neuron subcluster. **(A)** UMAP visualization of excitatory neuron subcluster cell scRNA-seq data from individual time points. Cells are colored by excitatory neuron subcluster assignment. **(B)** Top marker genes for each of the excitatory neuron subclusters.

**Figure S4.**
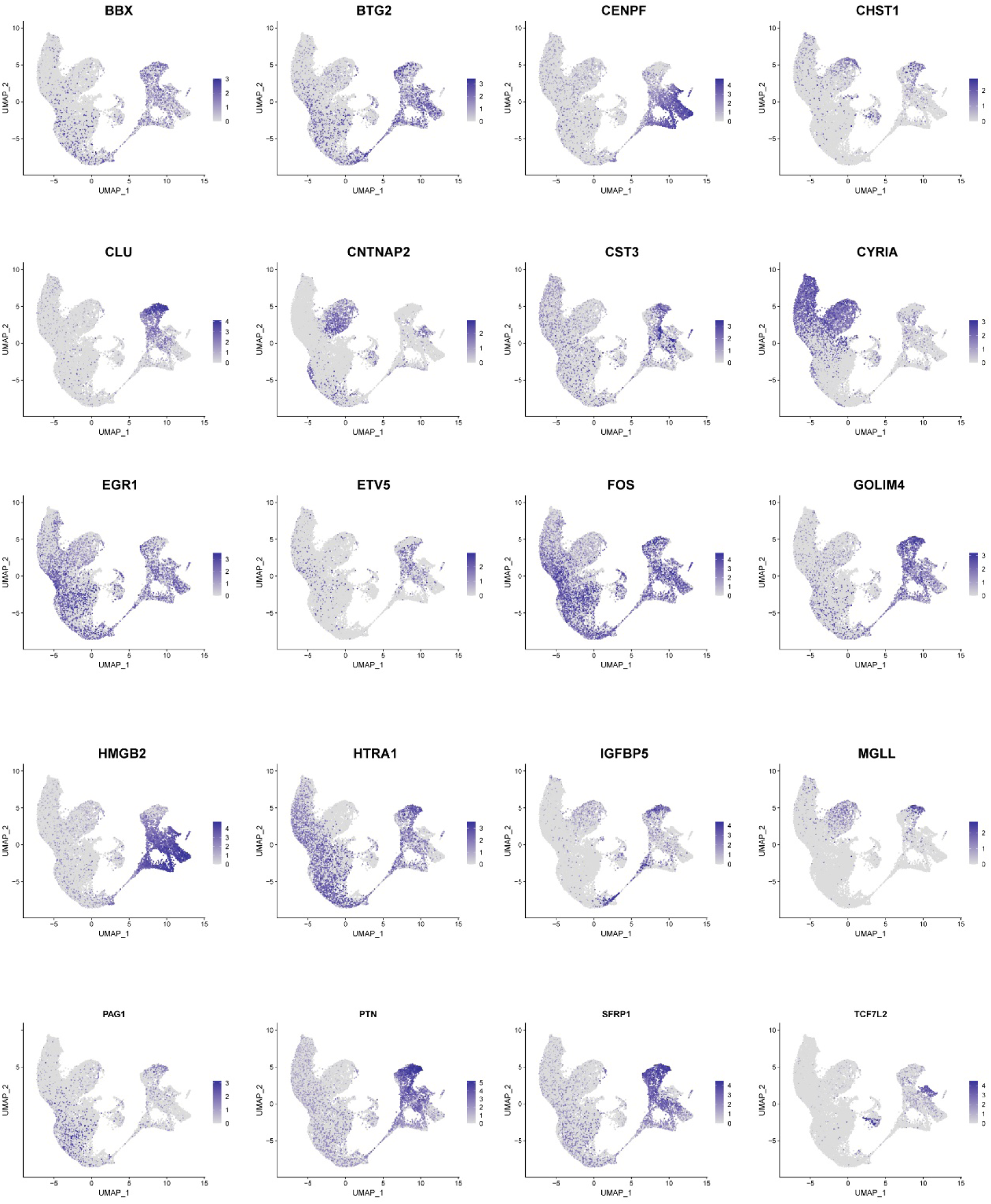
Stem and Excitatory neuron subclusters mapping genes.

**Figure S5.**
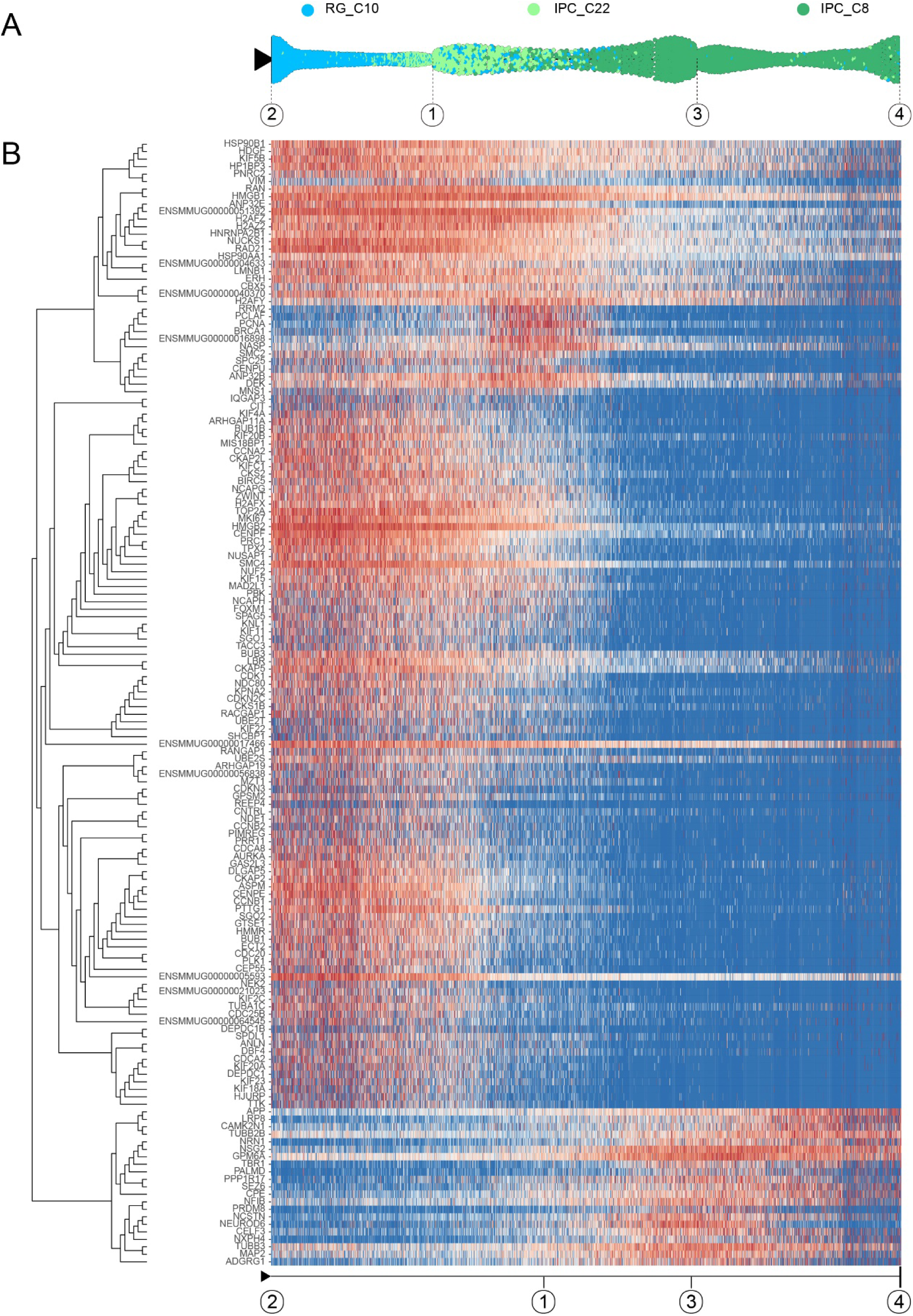
Developmental regulation of gene expression from RG to IPC. (**A**) Pseudotime analysis by Slingshot of EOMES-positive cells (C10-C22-C8). Dots: single cells; Cells are colored by their identity. **(B**) Heatmap shows the relative expression of top150 genes displaying significant changes along the pseudotime axis of RG to IPC (C10-C22-C8). The columns represent the cells being ordered along the pseudotime axis. The start point, endpoint, and important nodes of the Slingshot inference trajectory were marked by framed numbers. Framed number “2” was radial glial cells at the beginning of pseudo time (C10). Framed numbers”1” and “3” marked intermediate state nodes. Framed number “4” marked intermediate progenitor cell (C8).

**Figure S6.**
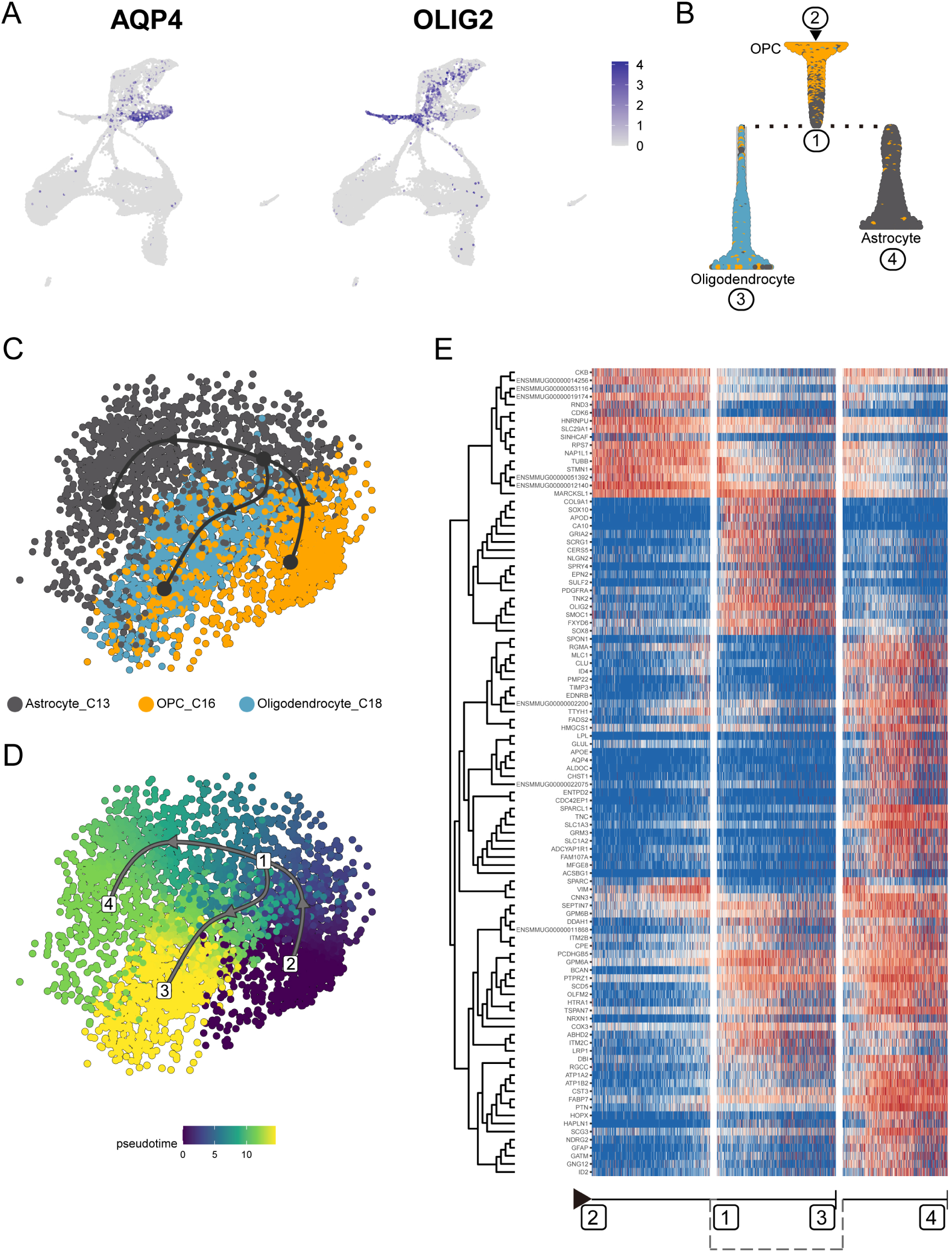
Transcriptional regulation of OPC differentiation into astrocytes and oligodendrocytes. (**A**) Feature plot of classic marker for astrocytes (AQP4) and oligodendrocytes (OLIG2). **(B)** Slingshot branching tree related to Slingshot pseudotime analysis in **(C)** and **(D)**. Root is OPC (C16), tips astrocytes (C13), and oligodendrocytes (C18). **(C** and **D)** Pseudotime analysis by Slingshot of glial cell lineage (C16→C13, C16→C12). The Slingshot result indicated the trajectories of lineages, and the arrows indicated directions of the pseudotime. Cells are colored according to their identity **(C)** and pesudotime (**D**). Dots: single cells; colors: cluster and subcluster identity. **(E)** Heatmap shows the relative expression of top150 genes displaying significant changes along the pseudotime axis of glial cell lineage (C16→C13, C16→C12). The columns represent the cells being ordered along the pseudotime axis.

**Figure S7.**
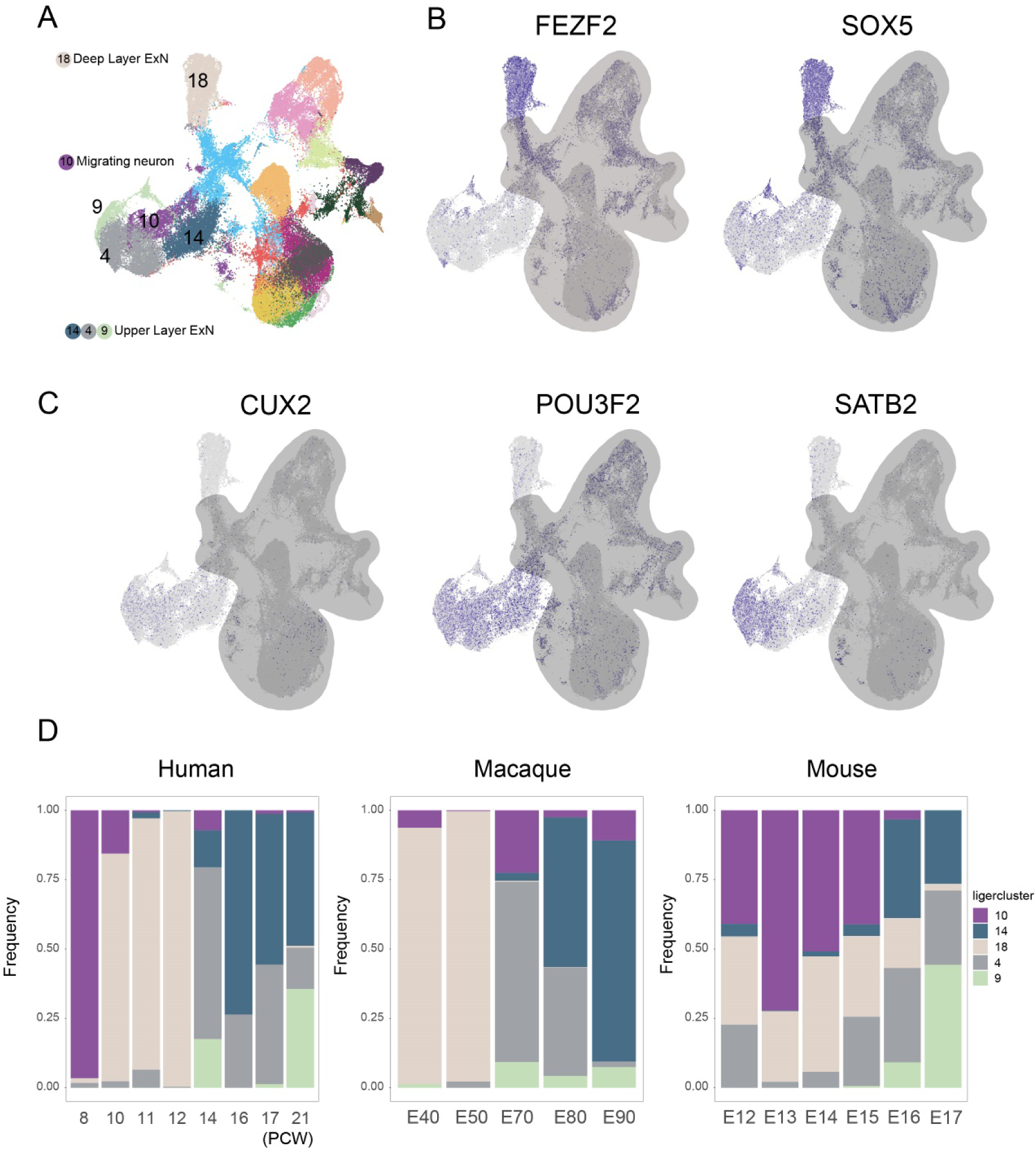
Upper layer and deep layer excitatory neuron proportion analysis among species. (**A**) Cell type annotation of the integrated dataset. **(B** and **C)** Feartureplot shows the expression of upper-layer marker genes *CUX2*, *POU3F2*, *SATB2*, and deep-layer marker genes *FEZF2* and *SOX5*. (**D**)Proportion analysis of excitatory neuron subclusters in human, macaque, and mouse datasets at different developmental time points.

**Figure S8.**
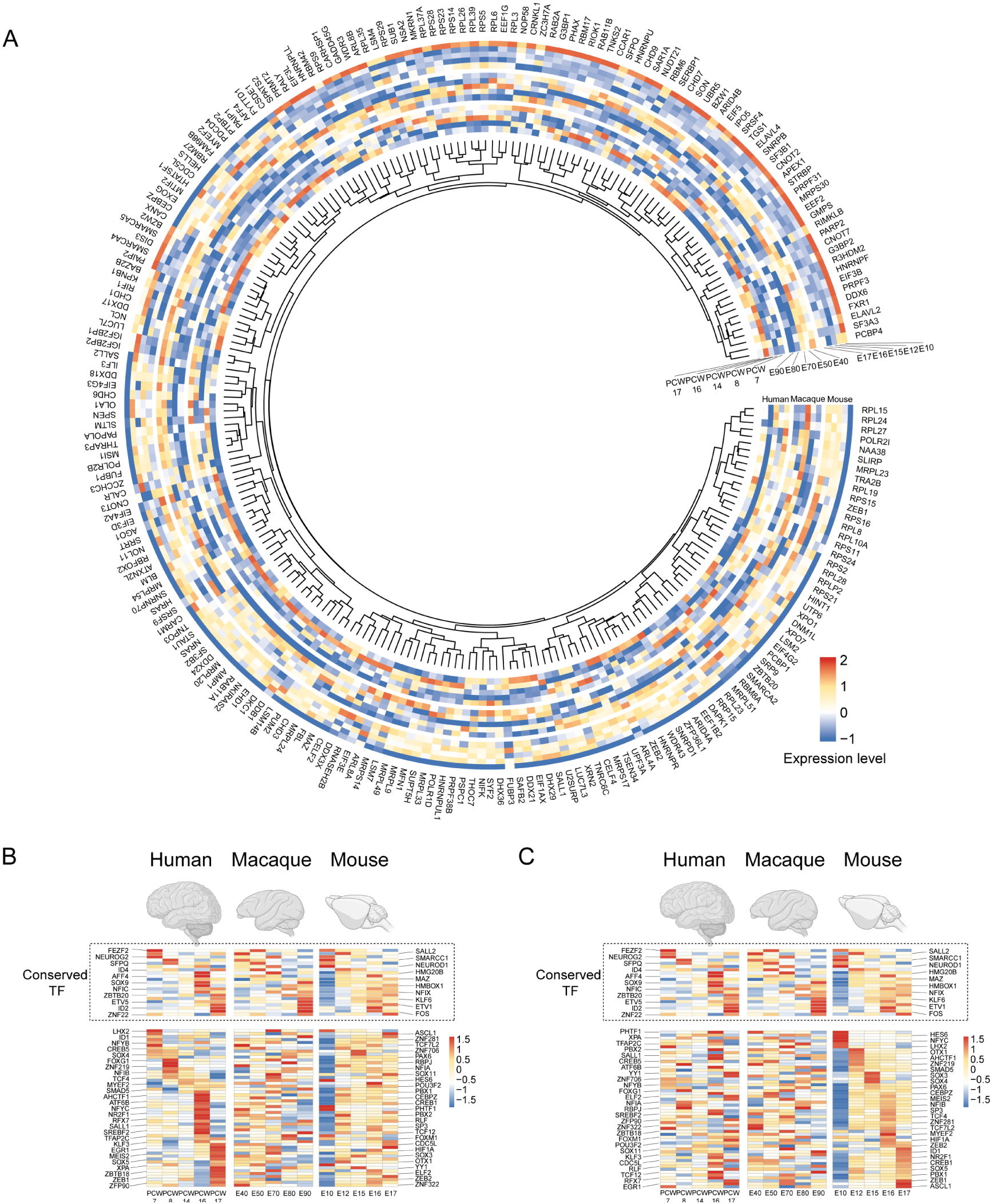
Temporal expression pattern of RNA binding protein and transcription factor genes in human, macaque, and mouse vRG. **(A)** Temporal expression heatmap of RNA binding protein genes in human, macaque, and mouse ventricular radial glia. **(B and C)** Temporal expression heatmap of Transcription factors genes in human, macaque and mouse ventricular radial glia sorted by human **(B)** and mouse **(C)** temporal pattern. (Note: each line is the same homologous gene).

**Dataset S1**. (separate file)

Marker list of 28 cell clusters.

**Dataset S2**. (separate file)

Gene expression importance across the vRG to oRG (C10-C14-C12) lineage trajectory related to Fig. 3E.( vRG: milestone2; oRG: milestone4)

**Dataset S3**. (separate file)

Gene expression importance across the RG to deep-layer neurons and upper-layer.

**Dataset S4**. (separate file)

Gene expression importance across the vRG to IPC(C10-C22-C8) trajectory related to fig. S5B.

**Dataset S5**. (separate file)

Gene expression importance across the OPC differentiation into astrocytes and oligodendrocytes lineage trajectory related to fig. S6E.

**Dataset S6**. (separate file)

The original data of normalized gene expression matrix in human, macaque, and mouse vRG related to fig.6A.

**Dataset S7**. (separate file)

Regulon specificity score for each timepoint in human, macaque, and mouse vRG related to fig. 6 C to H.

## Notes

### Competing Interest Statement

The authors have declared no competing interest.

### Summary of Updates

Figures 1 and 2 were revised to improve the cell definition; Figure 5 has been revised to improve the species comparative analysis section; Figure 1, figure 3, and figure 4 were revised to improve the visualization of the results. Supplemental files and datasets updated.

